# Molecular and functional profiling unravels targetable vulnerabilities in colorectal cancer

**DOI:** 10.1101/2024.04.17.589725

**Authors:** Efstathios-Iason Vlachavas, Konstantinos Voutetakis, Vivian Kosmidou, Spyridon Tsikalakis, Spyridon Roditis, Konstantinos Pateas, Ryangguk Kim, Kymberleigh Pagel, Stephan Wolf, Gregor Warsow, Antonia Dimitrakopoulou-Strauss, Georgios N Zografos, Alexander Pintzas, Johannes Betge, Olga Papadodima, Stefan Wiemann

## Abstract

While colorectal cancer (CRC) patients with microsatellite instability (MSI) respond well to immunotherapy those with microsatellite stable (MSS) tumors rely on conventional chemotherapy, often with poor outcomes. Both types frequently carry mutations in *KRAS* or *BRAF* proto-oncogenes, rendering them more resistant to treatment. New therapeutic biomarkers and treatments remain a clinical need, especially for MSS tumors. We performed whole exome and RNA-Sequencing from 28 tumors of the Athens Comprehensive Cancer Center CRC cohort, and molecularly characterized colorectal cancer patients based on their MSI status, SNVs/CNAs, and pathway/transcription factor activities at the individual patient level. Variants were classified using a new computational score for integrative cancer variant annotation and prioritization. Complementing this molecular data with public multi-omics datasets, we identified activation of transforming growth factor beta (TGFβ) signaling to be stronger activated in the MSS patients whereas JAK-STAT and MAPK molecular cascades were activated specifically in MSI. We unraveled mechanisms consistently perturbed in the transcriptional and mutational circuits and identified RUNX transcription factors as putative novel targets. Assessing the immunogenicity of CRC tumors in the context of RAS/RAF mutations and MSI/MSS status revealed a critical impact *KRAS* mutations have on immunogenicity particularly in the MSS patient subgroup, with implications for diagnosis and treatment.

## Introduction

Colorectal cancer (CRC) is a highly heterogeneous disease associated with different molecular background and clinical management. Improved screening strategies, surgical techniques and advanced treatment protocols have led to better clinical outcomes of CRC patients in the last decades ^1, 2^. The recent development of covalent inhibitors specifically targeting the KRAS G12C variant have indicated significant progress in targeting even previously ‘undruggable’ onco-proteins ^3^. Despite these advances, CRC has remained the second and third leading cause of cancer-related death in the US and worldwide, respectively ^4, 5^. Furthermore, recent epidemiological data has indicated a concerning trend of increased CRC incidence particularly in younger individuals below the age of 55, coupled with more advanced disease stages ^4^. The response to treatment is highly variable and not durable in the majority of cases. This dismal clinical perspective is impacted by various factors, including tissue tropism effects that allele-specific *RAS* mutants have in combination with other actionable alterations ^6^, clonal heterogeneity ^7^, phenotypic plasticity ^8^, location of tumor ^9^, and the tumor microenvironment ^10^. Collectively, these findings highlight the need for novel therapies focused specifically on the personalized treatment of CRC. The advent of next-generation sequencing has led to substantial progress in molecular characterization of the disease. Alterations in a number of genes with critical roles in CRC have been identified, such as in *APC, BRAF, EGFR, NRAS, KRAS* and *PIK3CA* ^11, 12^. The resulting aberrant activation of signaling pathways may be exploited in precision therapies. Recent accomplishments include combinatorial strategies targeting co-activated pathways in various types of unresectable CRC such as BRAF V600E mutated ^1^ and *HER2*-positive carcinomas ^2^.

The microsatellite instability (MSI) status is among the most informative parameters determining treatment decision towards personalized medicine in CRC ^13^, and in other tumor entities ^14^. According to the MSI status, CRC is categorized into three types: high microsatellite instable (MSI-H), low microsatellite instable (MSI-L) and microsatellite stabile (MSS) ^13^. MSI-L and MSS tumors are commonly grouped into one type ^15^. Accordingly, we use MSS for MSI-L/MSS and MSI for MSI-H tumors throughout. MSI-status accounts for ∼15% of CRC cases and is associated with sporadic or hereditary disease etiologies. While the former is associated with *BRAF* mutations ^16^ and *MLH1* promoter methylation ^17^, the latter has been associated with mutations mostly in mismatch repair genes *MLH1*, *MSH2*, *MSH6* as well as *PMS2* and is classified as Lynch syndrome ^18^. MSI tumors have a high mutational burden, resulting in the expression of neo-antigens that may be recognized by the immune system. Differences in the immune microenvironment thus distinguish MSI and MSS CRCs, and have immediate therapeutic impact: MSI CRC tumors mostly respond to immunotherapy, which has revolutionized clinical practice and outcome of respective patients ^19^.

In contrast, MSS CRC patients commonly do not benefit from immunotherapies and have mostly limited targeted treatment options, wherein the most informative molecular distinction involves the *RAS* mutation status ^20^. Despite the unmet therapeutic demand particularly for MSS patients, only few studies have investigated the multi-layered differences in MSI/MSS tumors, their downstream signaling effects, and their interrelation with *RAS*/*RAF* mutations in regard to immunogenicity and pathway activation ^21, 22^. Hence, by leveraging a prospective patient cohort of the Athens Comprehensive Cancer Center (ACCC – https://www.accc.gr/index.html) and two public cancer genomics datasets, we aspired to elucidate the molecular complexity of microsatellite (in)stability status in CRC and to uncover biological mechanisms that might be exploited to enhance therapeutic options particularly for patients with the MSS type.

## Results

### The mutational landscapes of MSS and MSI CRCs

To elucidate the landscape of somatic mutations in CRC, whole exome sequencing (WES) as well as RNA sequencing (RNAseq) was performed in cancerous as well as in adjacent normal tissues from 28 patients of a prospective patient cohort of the Athens Comprehensive Cancer Center (ACCC) (Supplementary Table 1). Since microsatellite instability is one of the major clinical factors determining therapeutic management and prognosis of CRC patients, we first wanted to infer the microsatellite stability status of every patient using RNAseq data. To this end, we initially validated the robustness of PreMSIm ^23^ and predicted the MSI status in 434 patients of the TCGA PanCan COAD/READ ^11^ as well as in 79 patients of the CPTAC ^24^ cohorts. PreMSIm estimates were highly concordant with MSI-MANTIS predictions from TCGA (concordance rate: 0.988, Supplementary Table 2), and with the annotated MSI-status of patients in CPTAC (concordance rate: 0.987, Supplementary Table 3). We then characterized the ACCC CRC samples applying PreMSIm and inferred five tumors as MSI and 23 as MSS (Supplementary Table 4).

The exomes of ACCC CRC samples were sequenced at an average coverage of >300x (range: 195 to 375). In total, 25,913 unique protein-altering somatic variants were identified in 11,492 genes, including 21,917 missense, 1495 stop_gained, 423 splice site, 1871 frameshift and 162 inframe InDel mutations (Supplementary Table 5). The number of somatic variants varied greatly among ACCC patients as the tumor mutational burden (TMB) ranged from 0.028 to 375.994 protein-altering mutations per Megabase (Supplementary Table 4). The median was 108 nonsynonymous SNVs in MSS tumors while that in MSI tumors was 1257. Five patients carried hot-spot mutations in the G12 codon of KRAS [G12D (n=3), G12V (n=1), G12S (n=1)], another five patients carried hot-spot mutations in BRAF [V600E (n=4), K601E (n=1)], while concomitant mutations in KRAS [A146V] and BRAF [D594N] were observed in one ACCC-patient. Another patient carried an HRAS [A146T] mutation co-occurring with two non-hotspot mutations in BRAF [L505F and L577I]. The HRAS [A146T] substitution has been found as a germline variant in Costello’s syndrome ^25^ and as a somatic mutation in a few cases of melanoma and other skin cancers (https://www.cbioportal.org, GENIE Cohort v14.1-public, accessed on 2023/12/04). BRAF [L505F] is a class II mutation ^26^ and has been shown to confer a RAS-independent constitutively activated state in the kinase domain ^27^. This and BRAF [L505H] have been reported in a small number of tumor entities including melanoma, prostate and pancreatic cancer in cbioportal. BRAF [L505H] was shown to be activating mutation conferring resistance to vemurafenib ^28^, and is characterized as oncogenic in OncoKB ^29^ (https://www.oncokb.org/gene/, accessed 2023/12/13). The BRAF [L577I] mutation has been reported in cutaneous melanoma and is characterized as variant with ‘Unknown Oncogenic Effect’ in OncoKB.

In order to prioritize somatic mutations according to their putative oncogenicity, we next applied our SVRACAS variant scoring pipeline (https://github.com/Jasonmbg/Simple.-Variant-Ranking-Annotation-CAncer-Score, accessed 2022/10/24) to the 25,913 variants, leaving 4307 prioritized variants (score ≥0.5) in 2969 genes (Supplementary Table 5). Amongst these, 54 matched with 72 CRC driver genes annotated IntOgen ^30^ (Fig. 1). The top frequently mutated genes in the ACCC tumors were *APC* and *TP53* (Fig 1) and the presence of mostly loss-of-function mutations was in line with previous observations ^11, 31^. Next, we revealed differences in the 54 mutated genes that had matched the IntOgen set, between the MSI and MSS CRC tumors of the ACCC cohort regarding somatic SNVs, InDels and copy number alterations (CNAs). On the one hand, MSS tumors exhibited a higher percentage of CNAs: Deletions of *SMAD2*, *SMAD4* and *MAP2K4*, and gains in *RNF6*, *BRCA2, NBEA* (chr 13q12-13) as well as in *GNAS* (chr 20q13.32) were only observed in MSS tumors. On the other hand, frameshift and missense variants were more frequent (*RNF43*, *ACVR2A*, *TGFBR2*) or exclusive (*BCL9L*, *PTPRC*) in the MSI cluster.

**Fig. 1.**
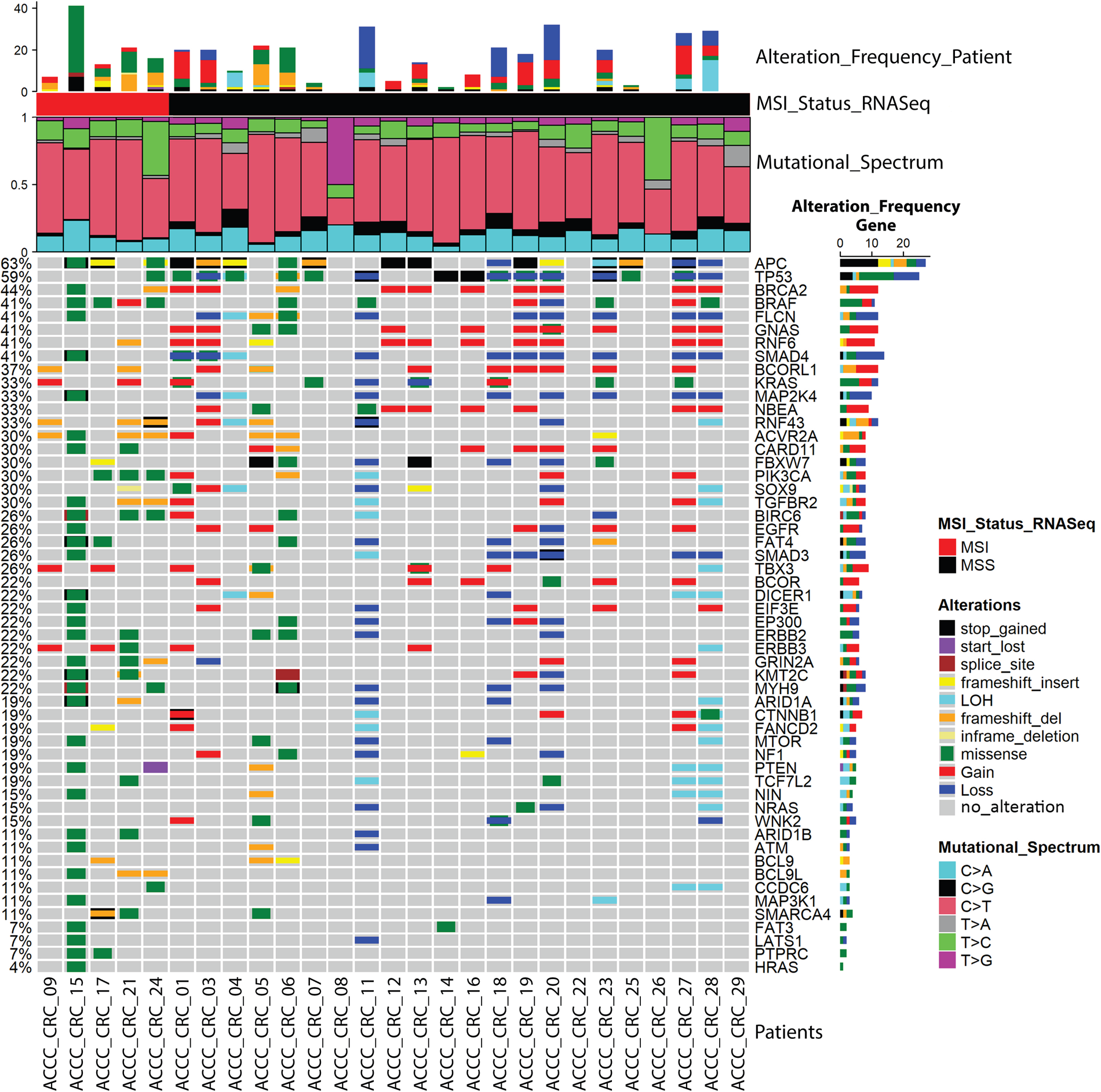
Mutational landscape in the tumors of the ACCC CRC cohort. Oncoplot presenting scored genes common with the IntoGen database for 27 ACCC-CRC patients, which were grouped into MSI and MSS subtypes (MSI_Status). Each column represents a profiled patient and each row a gene of interest. The alteration frequencies of 54 genes (SNVs, InDels and CNVs) are shown for each patient and for each gene on the top and on the right barplots, respectively. Genes are sorted based on their total mutation frequency (depicted on the left). Colored cells indicate respective types of alterations explained on the right, while gray cells indicate lack of mutations in respective tumors. The mutational spectrum for each patient showing the relative contribution of each of the six base substitution types is presented below the top bar plot as a stacked bar chart, using the Mutational Patterns R package ^116^ (version 3.4.1).

Mutational analysis implementing the COSMIC SBS signatures ^32^ revealed distinct patterns between the MSI and MSS tumors (Supplementary Fig. 1). The majority of the MSI tumors were characterized by SBS6, SBS26, and SBS20 signatures, all related to defective mismatch repair and, thus, microsatellite instability. One sample (ACCC_CRC_15) was further characterized by signatures 10a and 10b, which are related with mutations in *POLE*. Mutations in *POLE* are associated with a very high TMB, which was indeed observed in this patient (375.994 mutations/Mb). On the contrary, MSS samples were mainly enriched in the SBS1 and SBS5 signatures, which are related to age, stem-cell division and “clock-like” processes, and are in line with a general low TMB in MSS tumors. However, the DNA mismatch repair-associated signatures SBS6 and SBS15 were evident also in two MSS cases (ACCC_CRC_05 & ACCC_CRC_06, respectively), possibly related to their high TMB. Patients ACCC_CRC_07 and ACCC_CRC_18 were enriched with the unknown etiology signatures SBS93 and SBS94, which have been identified in gastric and colorectal samples, respectively. Taken together, these findings are in line with previous studies recapitulating the mutational landscapes of CRC, where microsatellite instability is associated with a low frequency of CNAs and a high TMB ^33, 34^.

### Mapping mutational landscapes to functional networks

To extend the analysis of mutational profiles in MSS and MSI CRC tumors, we utilized the public datasets of the TCGA PanCan COAD/READ ^11^ and the CPTAC ^24^ cohorts. Complete datasets were available for 434 patients (392 MSS, 42 MSI) from TCGA (Supplementary Table 2), and 79 patients (65 MSS, 14 MSI) from CPTAC (Supplementary Table 3). Highly scoring variants from SVRACAS annotation (score ≥0.5) were identified in 7287 and 2588 genes in the TCGA and CPTAC datasets, respectively (Supplementary Tables 6 and 7). Of those, only three genes had higher mutation frequencies in MSS compared to MSI tumors: *APC* and *KRAS* genes were found more frequently mutated in MSS in both public datasets, whereas mutations in *TP53* were significantly enriched only in the TCGA PanCancer dataset. In contrast, 564 (TCGA) and 80 (CPTAC) genes had significantly higher mutation frequencies in MSI compared to MSS tumors, and 42 genes were shared in both datasets. To uncover molecular networks possibly related to microsatellite instability, we then performed functional enrichment analysis by mapping the genes differentially mutated (MSS vs. MSI) in TCGA and CPTAC to Reactome^35^. Eleven pathways were significantly enriched in MSI vs. MSS tumors in both, TCGA and CPTAC. These signaling pathways were related to chromatin modification and organization, growth factor, second messenger as well as Wnt signaling, and to RUNX1 transcriptional regulation (Fig. 2). In line with only three genes having higher mutational frequencies in MSS vs. MSI, no Reactome pathway was enriched in MSS tumors.

**Fig. 2.**
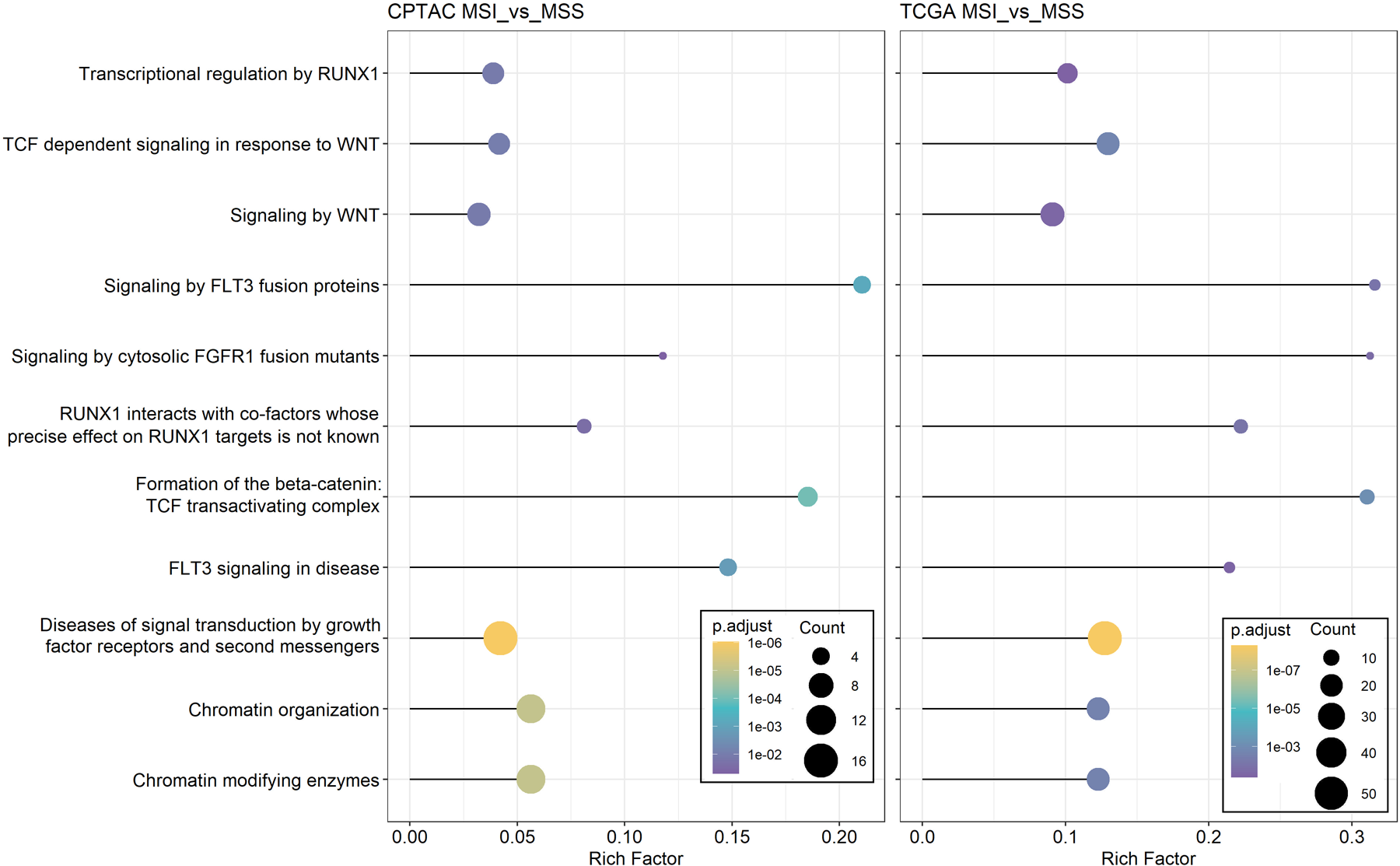
Lollipop plots of REACTOME pathways significantly enriched in MSI vs. MSS tumors of the TCGA and CPTAC cohorts. Functional enrichment analysis was performed using clusterProfiler ^104^ (v.4.2.2), and ggplot2 (v.3.4.3) was used for the visualization of enrichment results. P-values were adjusted for multiple comparisons (p.adjust) ^117^. Sizes of bullets represent the number of over-represented genes (Count) in the corresponding pathway and adjusted p-values (p.adjust) are color-coded. The Rich Factor value (x-axis) describes the relative ratio of the enriched genes in each pathway, divided by the total number of annotated genes in the respective set (i.e., over-representation analysis) ^118^.

Next, we mapped the 2969 mutated genes from the ACCC CRC dataset to the 11 enriched Reactome pathways. This identified 60 genes that were mutated with a higher frequency in ACCC MSI tumors and in at least one of the two other CRC datasets. Functional and physical associations were investigated by mapping these genes to the STRING database ^36^, identifying three protein interaction networks with 60 genes and 154 edges (Supplementary Fig. 2). The first network comprised of genes in angiogenesis as well as ERBB2, PI3K/AKT and PDGFR signaling. Network 2 was enriched in histone modifications, NOTCH1, TCF and Wnt/β-Catenin signaling, while network 3 was characterized by RUNX1 signaling, regulation of nucleotide-excision repair, double-strand break repair, and chromatin remodeling. The Reactome and STRING network topologies mirrored prior knowledge of MSI CRC thus further validating the ACCC CRC cohort and the classification of patients into MSS and MSI subtypes ^11, 12^.

### Molecular pathway and transcription factor activities are distinct in MSS vs. MSI mutated CRCs

Aiming to estimate the activity of cancer-relevant signaling pathways in MSS and MSI groups of the three CRC datasets (TCGA PanCan COAD/READ ^11^, CPTAC ^24^ and ACCC CRC), we next estimated the activity of 14 signaling pathways based on transcriptomic data using the PROGENy computational pipeline ^37^. The activity of only the TGFβ signaling pathway was significantly higher in MSS than MSI tumors in all three cohorts (Fig. 3A). This is in line with a recent report suggesting that TGFβ signaling in MSS tumors stimulates cancer associated fibroblasts and establishment of an immune-excluded tumor microenvironment ^38^. In contrast, six pathways were stronger activated in MSI tumors in all three cohorts (Fig. 3A). JAK-STAT signaling was the most activated pathway in MSI and has been implicated with positive immunotherapy outcomes ^39^. Then, kinase and pathway activity analysis was performed using the phospho-proteomics measurements from the CPTAC cohort. This suggested the cAMP-dependent protein kinase and TGFβ signaling pathways as more active in MSS tumors compared to MSI (Supplementary Fig. 3A), confirming the PROGENy results for TGFβ. Activation of Protein kinase A might be functionally related to gains and correlated expression of *GNAS* (r=0.45, p-value < 0.05) found specifically in MSS tumors also in the CPTAC cohort (Supplementary Fig. 3C, D). This pathway has been proposed as targetable driver mechanism in lung cancer ^40^. In contrast, several MAP kinases and PI3K signaling were significantly more activated in the MSI tumors (Supplementary Fig. 3A, B). The latter findings confirm that MSI tumors exhibit a higher activation of stress-response, inflammation/immune-modulation, and proliferation.

**Fig. 3.**
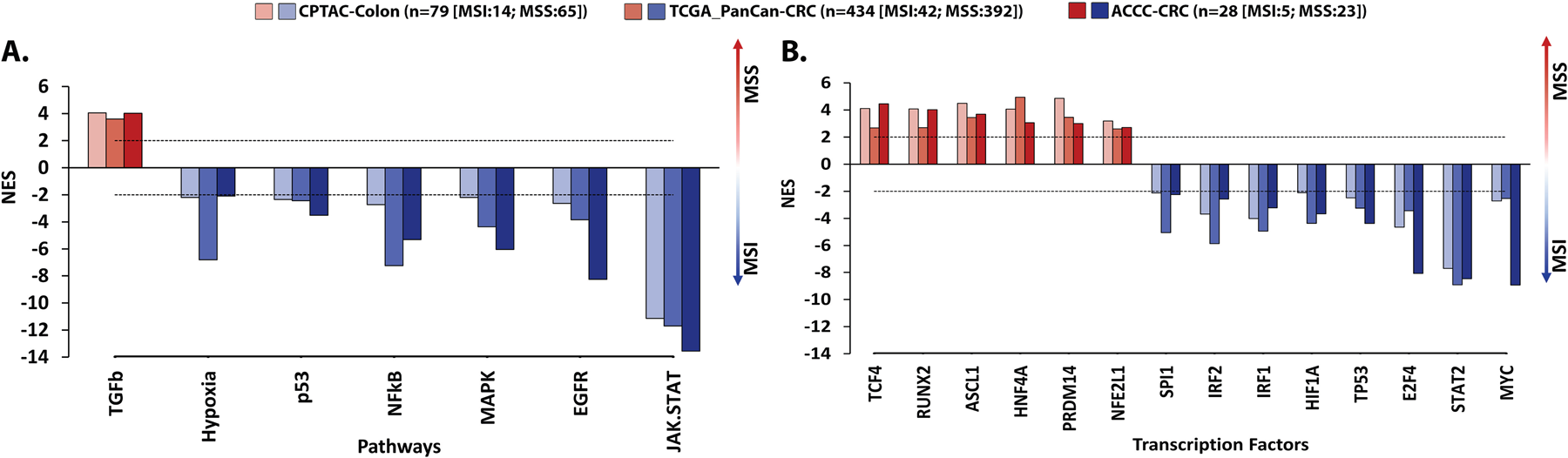
Significantly differentially activated pathways (A) and transcription factors (TFs) (B) between MSI and MSS CRC tumors in three CRC cohorts (CPTAC, TCGA and ACCC). Pathway and TF relative activities were estimated using PROGENy ^37^ and DorothEA ^109^ R packages, respectively. Barplot visualizes normalized Enrichment Scores (NES) with three different shades of red and blue color denoting significant pathway/TF activity levels of MSI and MSS tumors in CPTAC, TCGA, and ACCC CRC datasets, respectively. Horizontal dotted lines at +2/-2 values indicate the lower cut-off values for statistically significant activities of pathways (A) and TFs (B). Only pathways and TFs found significantly differentially activated in all three datasets are illustrated. The total numbers of samples characterizing each CRC group (MSI or MSS) are denoted in parentheses.

Next, we performed transcription factor (TF) activity analysis in MSS vs. MSI tumors and found 14 TFs that had consistently different activity profiles in all three datasets. Six transcription factors including RUNX2 (Runt-related transcription factor) were stronger activated in MSS CRCs compared to MSI in all datasets (Fig. 3B). RUNX transcription factors are downstream effectors of TGFβ, thus matching the results from pathway analysis. These TFs regulate basic cellular and developmental processes, stem cell biology, and tumorigenesis ^41^. In colorectal cancer, RUNX2 influences MAP kinase signaling via regulation of multiple RTKs, and its absence, can lead to resistance against MEK inhibitors along with its cofactor CBFB ^42^. MSI CRCs presented significantly higher activities for the MYC, STAT2, HIF1A, TP53, SPI1, IRF1 and IRF2 transcription factors (Fig. 3B). Some of these have previously been associated with CRC and potentially impact immunogenicity, proliferation, DNA repair mechanisms and cellular stress responses ^43^. Collectively, pathway and transcription factor activation-profiling point at elevated inflammatory processes in MSI tumors while tumors of the MSS type appear to be associated with stem-cell features as well as an immune-excluded microenvironment.

### Transcriptomic stratification and immune deconvolution of ACCC-CRC samples

We then estimated altered pathway and TF activities at the individual patient level to achieve a personalized view on the ACCC patients. To this end, we used the scaled gene-expression values of every patient as input and focused on the seven molecular pathways and 14 TFs, which we had found significantly altered in the three cohort-based comparison of MSS vs. MSI CRC tumors. The resulting estimated pathway and TF activities grouped the ACCC CRC patients into three patient clusters (Fig. 4A). To gain further insights into the underlying mechanisms that characterize the identified clusters, we performed single sample Gene Set Enrichment Analysis (ssGSEA) ^44^, immune deconvolution ^45^, and inference of Consensus Molecular Subtypes (CMS) ^46^.

**Fig. 4.**
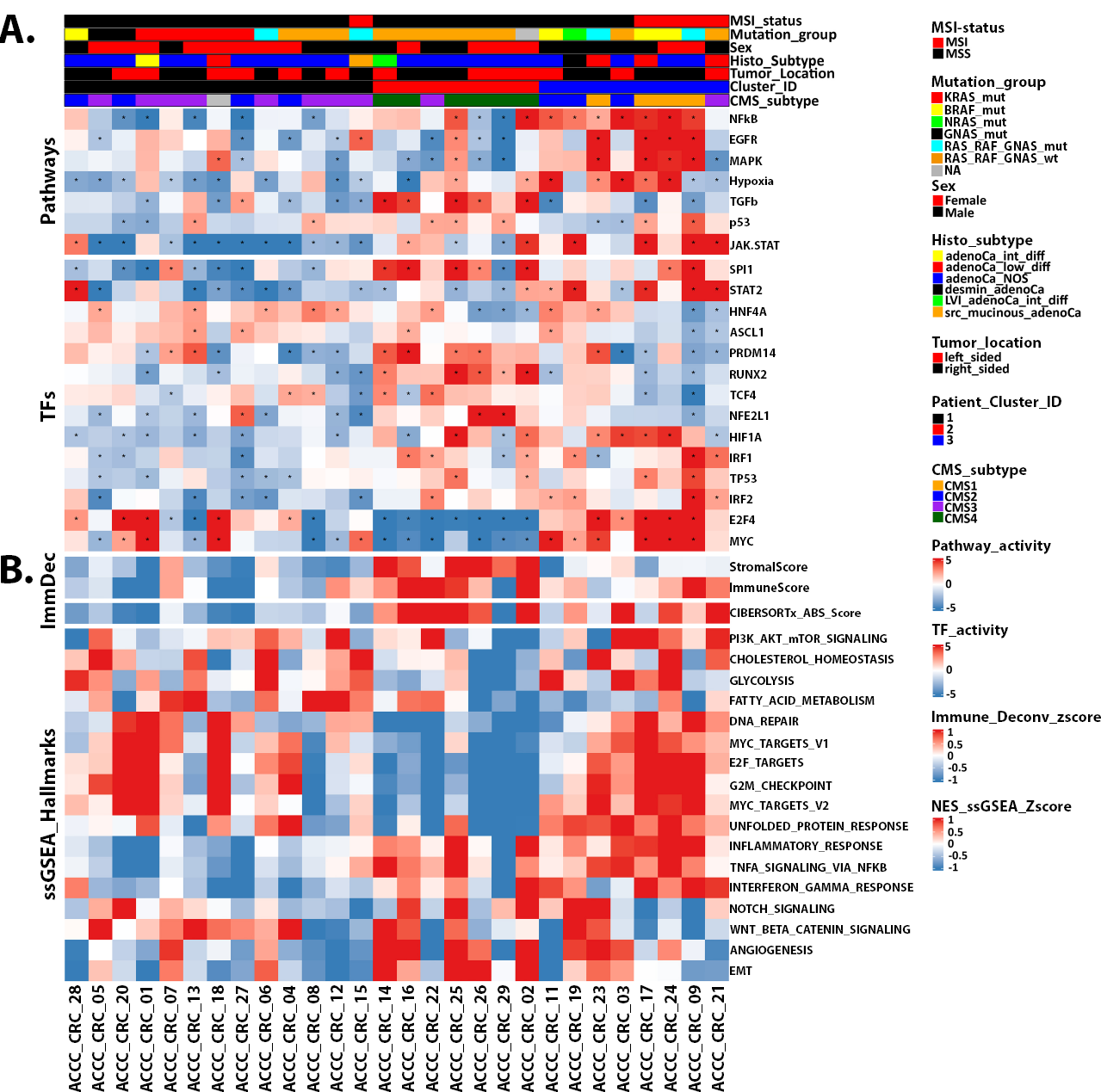
Clustering of ACCC-CRC patients based on integrative transcriptomic analysis. (A) Normalized enrichment scores (NES) of 7 pathway and 14 TF activities (compare Fig. 3) were used for unsupervised clustering on principal components (HCPC) with R packages FactoMineR ^114^ (version 2.8) and Factoshiny (http://factominer.free.fr/graphs/factoshiny.html, version 2.4), distributing ACCC patients into 3 patient clusters. Features were re-ordered within each Pathway and TF row sub-clusters using hierarchical clustering (“complete” method, “euclidean” distance), where samples were firstly grouped based on their cluster ID and subsequently based on their MSI status and mutational information. Tumors with co-occurring alterations in either in *RAS* and *RAF* or *GNAS* and *RAF* genes were assigned to the RAS_RAF_GNAS_mut group. Pathway and TF activities (NES) are indicated in shades of red (high) vs. blue (low). Asterisks in individual cells denote a significant activation of the respective pathway/TF in a patient (NES ≥2 or NES ≤-2). (B) Heatmap visualization of the normalized enrichment scores of hallmarks from MsigDb ^107^ per patient using ssGSEA ^44^. Re-ordering of rows was performed as described in (A). Hallmark pathway activities as well as immune or stromal scores (ImmDec: immune deconvolution), representing immune and stromal cell infiltration in respective tumors, are indicated in shades of red (high) vs. blue (low). Plots were created using the ComplexHeatmap R package ^115^.

The first patient cluster (Cluster_ID 1, n=13 patients) mostly included MSS (n=12) and CMS2 and CMS3 cancers (Fig. 4A). In line with MSS tumors in the TCGA and CPTAC (Fig. 3A), the tumors in cluster 1 were characterized by a low immunogenic behavior with sparse activation of the JAK-STAT and NFkB inflammatory pathways, no enrichment of immune related hallmarks, and flat immune infiltration scores (Fig. 4B). These findings are in line with the poor response MSS patients mostly have to immunotherapies. Patient ACCC_CRC_15, the only MSI sample in this cluster, also showed an overall trend of under-activation in the majority of estimated pathway and TF activities, whereas the EGFR signaling cascade and MYC TF were found significantly over-activated in this *POLE*-mutant tumor. This tumor had been diagnosed as serrated mucinous adenocarcinoma, which might explain the separate clustering from the other MSI tumors ^47^. The second cluster (Cluster_ID 2, n=7) contained only MSS and mostly CMS4 samples (n=6) which were characterized by an overall significant activation of the TGFβ signaling pathway, expression of genes associated with epithelial to mesenchymal transition (EMT) and angiogenesis as well as activation for the SPI1, RUNX2 and PRDM14. Almost all samples in the second cluster exhibited low activity of E2F4 and MYC, accompanied with reduced cell cycle activity and unfolded protein response (Fig. 4A, B). This suggests that tumors in the second cluster are characterized by an overall anti-proliferative capacity and a more invasive phenotype, which could be related to the molecular and clinical characteristics of the mesenchymal subtype (CMS4) ^48^. The third cluster (Cluster_ID 3, n=8) comprised four MSI and four MSS samples. It was mainly composed of CMS1 and CMS2 tumors with the exception of sample ACCC_CRC_21 that was predicted as CMS3. Tumors in cluster 3 were characterized by a relatively stronger activation of inflammatory (JAK-STAT, NFkB) and mitogenic (EGFR and MAPK) signaling pathways, compared to patients in the other clusters. Moreover, MYC and E2F4 seemed to be strongly activated. Conversely, a group of TFs, mainly associated with chromatin remodeling (PRDM14, ASCL1, RUNX2) and developmental processes (HNF4A, TCF4) displayed a reduced activity level compared to the other two clusters. This was most evident in the MSI samples of the third cluster, while MSS samples in this cluster showed a more variable activation pattern. Collectively, patients forming the third cluster reflected a more immune-reactive and less aggressive molecular profile.

The MSS tumors showed an overall higher degree of heterogeneity, which was associated with the prevalence of *KRAS* and *BRAF* mutations. *KRAS*-mutated MSS samples showed a trend towards having lower activation of immune response pathways, i.e. having significantly lower activities in JAK-STAT and NFkB pathways relative to the RAS_RAF_GNAS_wt MSS samples in the second patient cluster (Fig. 4A). This was corroborated by ssGSEA findings and immune deconvolution estimates (Fig. 4B), suggesting that *KRAS*-mutant MSS tumors in the first patient cluster were less immune reactive compared to the MSS tumors in the other clusters. These findings are in line with the observed low activity of IRF1 and IRF2, regulators of the interferon response ^49^, in the majority of the *KRAS*-mutant MSS samples of the first cluster (Fig. 4A). This is consistent with previous studies, which have identified a mutant *KRAS*-*IRF2* molecular cascade, promoting an immunosuppressive environment in CRC and fostering immunotherapy resistance ^50^. Conversely, the transcriptomic circuits of MSS CRC tumors in the third patient cluster showed greater similarity to the respective circuits in the MSI tumors within that cluster, compared to the MSS samples in the other clusters. In summary, while the evaluated transcriptomic programs primarily differentiated MSS and MSI samples and corresponded with the prevalence of identified CMS subtypes, they also exposed differentially enriched patient-cluster and patient-specific gene expression networks.

### From Oncogenic Mutational Landscapes and Transcriptomic Features Towards Personalized Therapeutic Strategies

Having seen higher functional heterogeneity in MSS compared to MSI tumors and that only few pathways seemed to be consistently stronger activated in MSS tumors, we hypothesized that precision oncology approaches might be particularly useful for MSS CRC. Along these lines, we went out to unravel actionable alterations for individual ACCC CRC patients and interrelate them with their respective transcriptomic profiles. This should provide more holistic views on potential vulnerabilities and frame them as comprehensive personalized cancer maps, placing our special emphasis on MSS cases. For this purpose, the SVRACAS prioritized somatic variants in each patient were annotated with clinical actionability-information from the OncoKB precision oncology database ^29^. This analysis prioritized 34 actionable variants in 20 genes, reflecting these variants as predictive biomarkers of drug sensitivity or resistance to particular targeted therapies (Fig. 5A).

**Fig. 5.**
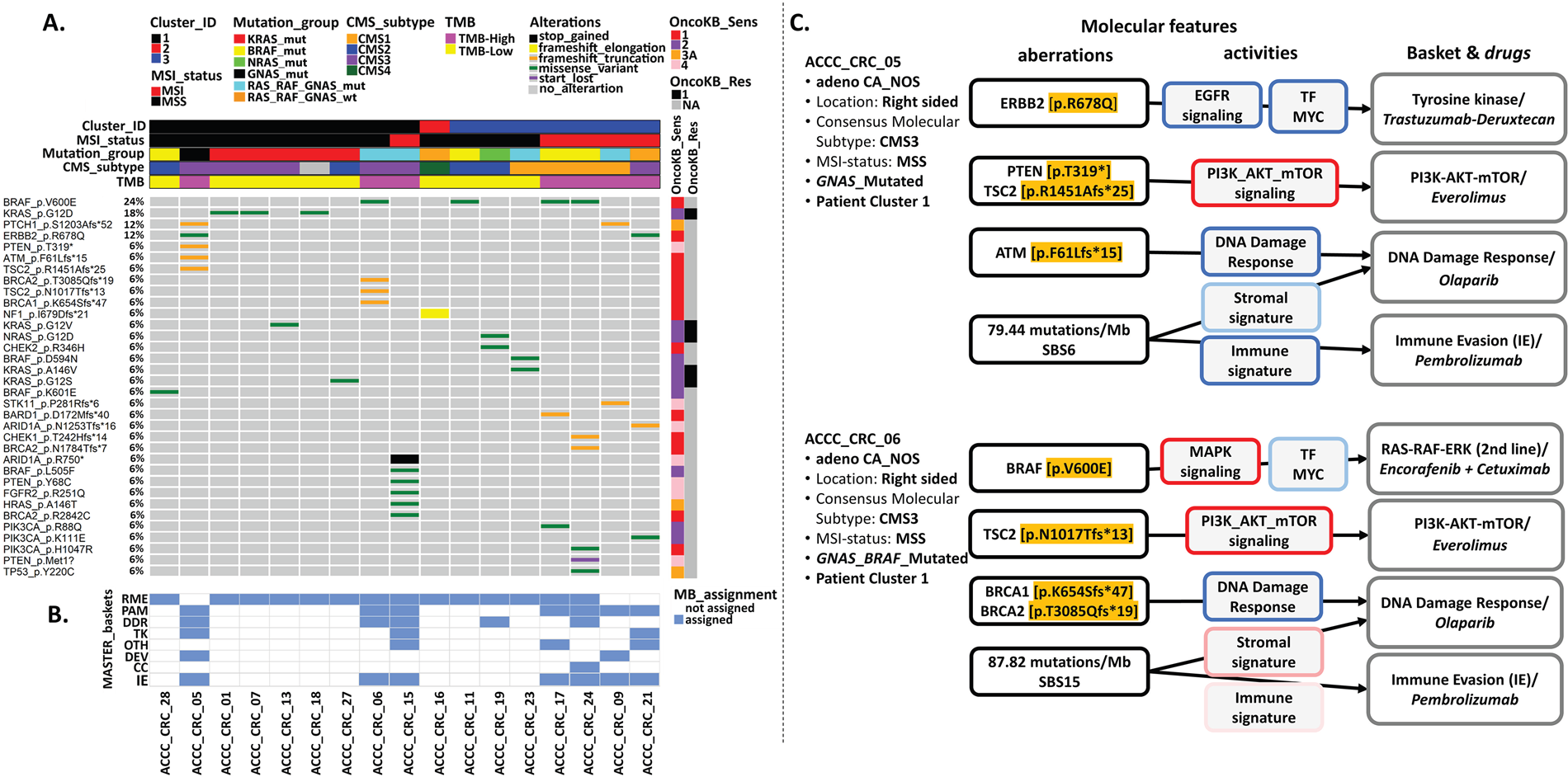
SVRACAS prioritized somatic mutations, pathway/TF activities, and immune profiles stratify ACCC CRC patients to MASTER baskets and personalized therapeutic interventions. (A). Oncoplot with actionable variants (SVRACAS score ≥0.5) and annotated in the context of sensitivity (OncoKB_Sens 1-4) or resistance (OncoKB_Res R1-2) to a targeted therapy (https://www.oncokb.org/levels) in any tumor entity. Columns correspond to patients and rows denote genes and protein-level types of alterations. Types of alterations are indicated in color, while wild type condition is indicated in gray. Rows were sorted by the percentage of occurring alterations, whereas columns/samples were firstly grouped based on patient cluster ID, then by microsatellite instability status, and subsequently by selected driver genomic alterations (i.e., *KRAS*/*BRAF*/RAS_RAF_WT). TMB: Tumor mutational burden. Tumors with co-occurring alterations in either in *RAS* and *RAF* or *GNAS* and *RAF* genes were assigned to the RAS_RAF_GNAS_mut group. (B). Mapping of indicated patients to MASTER interventional baskets (MB_assignment in blue) based on mutated genes that are associated with respective baskets ^119^ (RME: RAF–MEK–ERK; PAM: PI3K–AKT– mTOR; DDR: DNA damage repair; TK: tyrosine kinases; OTH: other; DEV: developmental regulation; CC: cell cycle). Mapping to the immune evasion (IE) basket was based on the presence of MSI and/or high TMB. MB_Assignment in white: not assigned. (C). Molecular rationale for suggested personalized therapeutic interventions for two ACCC_CRC patients. The first series of boxes (aberrations) illustrates actionable variants (in orange) and Tumor Mutational Burden (TMB) as well as MSI-related COSMIC SBS signatures ^32^. The second series (activities) presents perturbed molecular pathways, TF-MYC, and stromal and immune signatures, with boxes framed in shades of red or blue to indicate either high or low activity levels and high or low scores, respectively. The final series (Baskets & drugs) shows interventional baskets ^119^, with suggested drugs, based on a comprehensive analysis of actionable genomic variants, molecular pathway/TF activities, and immune profiles.

Next, we assigned the ACCC CRC patients to seven MASTER interventional baskets ^51^, based on the mutational profiles the respective tumors presented with. The MSI-status and TMB were regarded for assignments to the immune evasion (IE) basket. Seventeen patients of the ACCC cohort (∼63%) were stratified into at least one interventional basket (Fig. 5B). Tumors in patient cluster 1 were mainly characterized by overrepresentation of the RAS-MEK-ERK (RME, 8 out of 9 patients), the PI3K-AKT-mTOR (PAM, 3 out of 9 patients), and the immune evasion (IE, 3 out of 9 patients) baskets. An interventional basket (i.e., RME) could be assigned to just one (ACCC_CRC_16) out of seven MSS patients forming cluster 2, in line with the rather low TMB in the tumors forming this cluster. Cluster 3 was mostly represented by RME (5 out of 7 patients) and PAM as well as IE (4 out of 7 patients each) baskets.

The five MSI– and two MSS-patients could qualify for treatment with immune checkpoint-inhibitors (e.g., pembrolizumab), the MSS-patients in the light of the high TMB in the tumors (Fig. 5B). Two MSI and another two MSS tumors carried BRAF V600E mutations and might benefit from treatment with encorafenib and cetuximab ^52^. Seven patients carried *KRAS* or *NRAS* mutations, rendering those patients incompatible with EGFR-antibody therapy ^53^. Inhibitors specific for the most common KRAS variant G12D, which was present in three cases, are currently in preclinical and clinical development (e.g. MRTX1133, NCT05737706). Tumors carrying druggable KRAS G12C mutations ^3^ were not present in our cohort. No specific treatments were clinically available for the patients carrying other RAS mutations, i.e., ACCC_CRC_13 (G12V) and ACCC_CRC_27 (G12S). However, also non-specific Pan-KRAS inhibitors are in development ^54^, which might be applied once having proven efficacious. Hence, molecular profiling identified potential treatment options for the majority of patients in the cohort and more than one interventional basket was mapped to eight patients, suggesting a combination of targetable driver mechanisms in the respective patients.

We finally went ahead with two patients of the MSS type to prove the utility of our approach towards suggesting potential vulnerabilities for tailored therapeutic interventions. Patients ACCC_CRC_05 and ACCC_CRC_06 had presented with adenocarcinomas of the CMS3 subtype and had a right-sided location. Patient ACCC_CRC_05 (Fig. 5C) was characterized by an ERBB2 p.R678Q mutation, which has a modest biological rationale for treatment with trastuzumab-deruxtecan ^55^ (NCT evidence level m4). Furthermore, this patient carried an ATM p.F61Lfs*15 mutation, which has been found in few, mostly colorectal tumors (CosmicID: COSV53735143, accessed 2023/12/14). While preclinical evidence supports PARP inhibition ^56^ (NCT evidence level m3) this was recently shown (NCT02693535) to be ineffective in *ATM*-mutated CRC ^57^ thus not giving it high priority. In patient ACCC_CRC_05, we identified *PTEN* p.T319* and TSC2 p.R1451Afs.25 mutations, both of which have biological justification and preclinical evidence ^58^ suggesting treatment with mTOR inhibitors (NCT evidence level m3-m4). Tumor gene expression analysis for this patient demonstrated an over-activation of PI3K/AKT/mTOR signaling. The data further suggested an under-activation in the EGFR and JAK-STAT signaling pathways, which might indicate a predominantly PI3K/mTOR-driven cancer phenotype for this patient, further supporting the rationale for mTOR inhibition.

MSS patient ACCC_CRC_06 carried several potentially druggable alterations (Fig. 5C). The BRAF V600E mutation in this tumor could be targeted with Encorafenib and Cetuximab as an approved combination for second line targeted therapy after chemotherapy ^1^. BRCA1 p.K654Sfs*47 and BRCA2 p.T3085Qfs*19 mutations in this patient suggested PARP inhibition as a potential option. Tumor transcriptomic analysis revealed an over-activation of the PI3K/AKT/mTOR and under-activation of the DNA repair and JAK/STAT molecular circuits. A frameshift mutation in *TSC2* (p.N1017Tfs*13) would suggest mTOR inhibition as therapeutic strategy ^59^. However, in contrast to patient ACCC_CRC_05, we noticed a trend towards activation of the MAPK signaling pathway in patient ACCC_CRC_06, which might be associated with the presence of the BRAF V600E mutation. Thus, the value of targeting mTOR in the context of an activating BRAF mutation seems questionable ^60^.

Patients ACCC_CRC_05 & _06 both presented with high TMB, which has been observed in MSS tumors before ^31^, and mutational signatures associated with DNA mismatch repair (SBS6 and SBS15 in patients ACCC_CRC_05 and ACCC_CRC_06, respectively). While the TMB might indeed indicate application of immunotherapies, the lack of mutations in *POLE* ^61^, and the modest stromal and immune signatures in these tumors were not supportive of this (Fig. 4), suggesting immune checkpoint-inhibition as an only hypothetical treatment option ^62^. Yet, our analysis pointed at rationale treatment options for these two as well as for most other patients in the ACCC CRC cohort. Collectively, our study underlines the significant value of incorporating various molecular and functional layers to support therapeutic decision-making.

## Discussion

Here, we set out to characterize the molecular and activity landscapes MSS and MSI colorectal cancer. By implementing an integrative computational workflow we analyzed tumors from 28 patients of a prospective CRC patient cohort of the Athens Comprehensive Cancer Center (ACCC), in combination with two large public multi-omics datasets. There, we explored the molecular profiles of MSS vs. MSI CRCs, their relation to *KRAS*/*BRAF* mutations, and pathway as well as transcription factor activities in the context of personalized medicine. Firstly, we observed novel associations between particular genes and their increased mutational frequencies specifically in MSI tumors. Mutations were enriched in genes that are involved in molecular pathways mainly related to transcriptional regulation cascades by RUNX1, chromatin modification mechanisms, and in signaling cascades that are related to Reactome ‘fusion mutants/proteins’ (i.e., FGFR1/FLT3). These findings resonate with research indicating the central role of RUNX1 in the malignant transformation process and metastasis of CRC through mechanisms, like the Wnt/β-catenin signaling pathway and EMT ^63, 64^. Fusion events involving FGFR have been considered as therapeutic targets in various solid tumors ^65^. In contrast, MSS tumors demonstrated a pronounced enrichment in CNAs. Deletions were recurrently found in *SMAD2*, *SMAD4*, and *MAP2K4*, which are pivotal in the TGFβ and JNK signaling pathways ^66, 67^, respectively. Gains were observed in chromosomes 13q12-13 and 20q13.32, which have been associated with early events in the development of colorectal cancer ^68^, also affecting *GNAS*. These CNAs exclusively presented in MSS, suggesting a unique MSS genomic signature with implications in MSS disease etiology and therapeutic targeting ^69^.

Secondly, our comprehensive multi-omics analysis indicated differentially regulated circuits at the transcriptional level. Immune related biological mechanisms, such as JAK-STAT and NF_κ_B signaling pathways, were found upregulated in MSI tumors, consistent with the higher ‘immunogenicity’ in the context of microsatellite instability ^70^. Conversely, the TGFβ pathway showed greater activation in the MSS tumors and has been associated with epithelial-to-mesenchymal transition (EMT), metastasis, and therapy resistance ^71, 72^. We further observed that the synchronous presence of *KRAS* mutations with microsatellite instability had a stronger effect on the immune profile of CRC tumors than mutations in *BRAF*. Hence, we identified context-specific functional associations that are likely related to the higher functional heterogeneity within MSS CRC tumors. Previous studies have emphasized the intricate role *KRAS* mutations play in shaping the tumor microenvironment (TME) of colorectal cancer. These mutations seem to foster an immunosuppressive environment, which is marked by a diminished anti-tumor immune response ^73^.

Next, we identified transcription factors with differential activity between MSS and MSI CRC tumors. MSI CRC tumors showed higher activities of MYC, STAT2, HIF1A, TP53, SPI1, IRF1, and IRF2. These factors are instrumental in various cellular processes, such as cell growth, response to stress, oxygen homeostasis, and immune modulation ^43^. Hence, therapeutic regulation of TFs might be a promising strategy for CRC therapy owing to their underlying role as key effectors of various signal transduction and tumor-associated immune responses ^74^. Distinct transcriptional profiles were observed in MSS CRCs, indicating the heightened activity of RUNX2 upstream of TGFβ signaling ^75^. Yan et al. revealed that RUNX2 is critical for the maintenance of the stem cell-like properties of CRC cells as well as for the promotion of CD44-induced EMT in CRC^76^. Collectively, from our integrative analysis we noticed the significant deregulation of specific members of the RUNX TF family at DNA (*RUNX1*) or mRNA (*RUNX2*) levels, supporting their emerging role as prognostic biomarkers in CRC ^77^.

Our comprehensive molecular tumor analysis thus corroborated prior findings related to the phenotypic characteristics of CMS subtypes, however, also suggests that the microsatellite status is not the only biomarker that could predict molecular and functional tumor characteristics. In our integrative clustering analysis, one cluster grouped MSI with MSS patients together. This could be explained, in part, with epigenetic reprogramming in tumor progression, playing a major role in the clonal evolution and tumorigenesis of CRC ^78^. Indeed, this aligns with a recent study proposing a novel molecular classification of CRC tumors based on single-cell sequencing data ^79^. This study identified a subgroup of MSS samples (iCMS3_MSS), which exhibited more similarity with the transcriptional and genomic levels of MSI CRC tumors (iCMS3_MSI), forming a novel epithelial subtype ^79^. This may indicate that the MSI status, even though clearly distinct at the genomic level, may be viewed as a continuum through different states and sub-conditions at the transcriptomic layer ^80^. The potential therapeutic consequences of this, however, remain to be uncovered.

Finally, we sought to provide a personalization of molecular findings, as the identification of targetable alterations is a primary focus of current clinical research, to facilitate personalized treatment strategies. Effective therapeutic options exist for MSI CRCs employing immune checkpoint inhibitors. In contrast, the management of MSS cancers still predominantly relies upon conventional chemotherapies, as the most frequently encountered mutations in MSS CRC affect non-druggable tumor suppressor genes, such as *APC* and *TP53*. Indeed, the prevalence of well-established targetable mutations with clinical evidence supporting therapeutic interventions is exceedingly low in CRC. These primarily encompass BRAF V600E mutations, which we observed also in our cohort and have proven more difficult to target in CRC compared, for example, to melanoma ^52^. Clinical treatment options are often limited even when mutations in potentially druggable drivers have been identified ^81^. Hence, the integration of expression data might prove beneficial in clarifying therapeutic decisions for patients with uncertain mutation-driven treatment options, which we illustrated with patients ACCC_CRC_05 and ACCC_CRC_06. Our integrated analysis suggested treatment options for these patients (PARP/mTOR inhibition) that were unknown when the patients were indeed in clinical treatment.

The added value of integrating molecular modalities to foster precision medicine was initially introduced with the WINTHER clinical study ^82^ (NCT01856296). WINTHER was the first trial to describe the combination of matched genomic and transcriptomic information in metastatic tumors, showcased the significance of integrated molecular profiling into improved therapeutic recommendations, and expanded the portfolio of putative therapeutic interventions ^82^. Likewise, the more recent MASTER/NCT clinical study has illustrated that the integration of WES/WGS with RNA sequencing can result in a clinical benefit and boost patient treatment stratification in patients with advanced cancer of rare cancer entities ^51^. Our findings support these concepts, while we placed our focus on MSS and MSI CRC, in the context of the mutation status in the *KRAS* and *BRAF* genes. Our study initially suffered from the relatively small sample size of the retrospective ACCC cohort, which impeded direct generalization of inferential statistics and related biological comparisons. To cope with this limitation, we took advantage of public data sets of the TCGA and CPTAC cohorts, constructed an integrative workflow, and comprehensively profiled the sequencing data from the ACCC and the public multi-omics cancer datasets. Collectively, this allowed us to perform a patient-centric analysis capitalizing on the available molecular information as a guide to therapeutic interventions. In conclusion, our approach integrates mutation ranking with immune deconvolution and pathway as well as transcription factor activity analysis, to suggest personalized treatment strategies. In the future, this will likely be complemented with proteomic information ^83^ and holds the promise for enhancing the biological rationale behind individualized therapeutic interventions even it does not constitute a therapeutic protocol per se. Yet, it serves as a paradigm that supports and augments the concept of tailored medicine in oncology.

## Methods

### Patients & clinical specimens

Colorectal cancer patients operated in the surgical department of Gennimatas General Hospital in Athens, which is part of the Athens Comprehensive Cancer Center (ACCC), participated in the study. The study was approved by the ethics committee of the Gennimatas Hospital (protocol number 27659). Written informed consent was obtained from all patients. Only patients (n=28) with histologically confirmed primary colorectal adenocarcinoma and one case of rectal cancer who had received radiotherapy before surgery, were enrolled in the study cohort. Specimens from each patient were obtained from both the primary tumor and corresponding normal mucosa, which was taken from the histologically negative margins (at least 5 cm away from the macroscopically defined cancerous area) to ensure the absence of cancer cells. Upon surgery, resected samples were placed in 5-mL RNA*later* (Ambion, Austin, TX, USA) to preserve RNA integrity.

### DNA and RNA extraction

DNA and RNA were isolated with the AllPrep DNA/RNA Mini Kit (Qiagen, Hilden, Germany) according to the manufacturer’s instructions. Quantification of nucleic acids was performed with a Qubit 2.0 Fluorometer (ThermoFisher Scientific, Waltham, MA, USA) using the Qubit 1X dsDNA BR Assay (ThermoFisher Scientific) and RNA BR Assay kits (ThermoFisher Scientific) for DNA and RNA, respectively. In total, 28 paired tumor and normal DNA samples were isolated as well as 28 matching tumor RNA samples.

### Whole Exome Sequencing (WES) and variant calling

DNA libraries were prepared from tumor and matched normal tissues of ACCC patients using Low Input Exome-Seq Human v6 (Agilent Technologies, Waldbronn, Germany) for exome capture and analyzed on a NovaSeq 6000 (Paired-End 150bp S2, Illumina, San Diego, CA, USA) in the DKFZ High-Throughput Sequencing Core Facility. The resulting raw sequencing reads (median: 123 million reads/sample; spread: 80-148 million reads/sample) were preprocessed based on the internal WES alignment workflow (AlignmentAndQCWorkflows: 1.2.73-1), as part of the DKFZ-ODCF OTP pipeline^84^. Briefly, after initial quality control using FastQC, raw reads were aligned to a hg19 customized version of the 1000 genomes reference genome (ftp://ftp.1000genomes.ebi.ac.uk/vol1/ftp/technical/reference/phase2_reference_assembly_sequence/hs37d5.fa.gz) (1KGRef_PhiX), using bwa-mem (version 0.7.15). PhiX-genome sequence was added to the reference as an additional contig to remove spike-in sequences used to identify potential cross-contaminations and sample swaps. Marking of duplicated reads was performed with Sambamba tools (https://github.com/biod/sambamba/releases, version 0.6.5) embedded in the DKFZ-ODCF Roddy pipeline (https://github.com/DKFZ-ODCF/AlignmentAndQCWorkflows).

High-confidence functional somatic single nucleotide variants (SNVs) and InDels (Insertions & Deletions) were identified using DKFZ-ODCF SNVCallingWorkflow (https://github.com/DKFZ-ODCF/SNVCallingWorkflow, version 1.2.166-1) and IndelCallingWorkflow (https://github.com/DKFZ-ODCF/IndelCallingWorkflow, version 1.2.177-0), respectively, based on the BAM files of each tumor and its paired control sample. Various quality control criteria were performed to exclude spurious or/and low quality calls (such as identification of low quality bases, variant allele fraction and presence of strand bias artifacts), finally assigning a confidence value to each variant. Sample ACCC_CRC_02 was disregarded in further analysis as COSMIC SBS signature analysis (v3.4-October 2023) flagged this as having a potential PCR/sequencing artifact-based strand bias (prevalence of C>A alterations indicated presence of COSMIC SBS52; https://cancer.sanger.ac.uk/signatures/sbs/sbs52/, accessed 2023/12/21). Finally, the ANNOVAR tool ^85^ (version February 2016) for variant annotation was utilized for the derivation of all functional mutations defined only by the exonic mutations including missense, nonsense, splice-site, and stop-loss alterations, by setting a confidence value of at least 8 out of 10. High-confidence InDels were selected using the Platypus tool ^86^ implemented in a custom workflow (https://github.com/DKFZ-ODCF/IndelCallingWorkflow, version 0.8.1.1).

### SVRACAS variant annotation and gene prioritization

To efficiently prioritize the resulting high-confidence somatic variants for each patient with the ultimate goal to identify informative oncogenic alterations, the SVRACAS custom variant scoring pipeline was applied (https://doi.org/10.5281/zenodo.5636747, accessed 2022/10/24). This integrative workflow consists of two major steps: variant annotation and prioritization of somatic functional alterations. Briefly, the SVRACAS ranking module integrates 15 biological resources using the open-source OpenCRAVAT/OakVar comprehensive workflow for cancer variant annotation and exploitation ^87^. The aggregated output has four evidence categories: “Pathogenicity evidence” (spanning variant effect prediction tools), a “Cancer evidence” layer (covering cancer variant interpretation knowledge bases), a “Clinical support” indication (nominating clinically relevant alterations), and “Expression information” indicating whether an affected gene is expressed (i.e., ≥ 10 read counts in at least 5 samples). An integrated scoring value was then assigned to each variant as a total SVRACAS score (https://github.com/Jasonmbg/Simple.-Variant-Ranking-Annotation-CAncer-Score). This holistic prioritization scheme ranges from 0 to 1, representing an average approximation of the four different types of evidence, based on single nucleotide changes and InDels that occur in the protein-coding space. Here, a cutoff value ≥ 0.5 was used to include only variants annotated in at least two of the four evidence categories. Overall, the selection of individual relative cutoffs and weighted importance of different sources of evidence was based on proposed standard operating procedures that built on guidelines from cancer consortia such as VICC ^88^ and ComPerMed ^89^. Protein-coding genes were then mapped to the MANE Select Set (version 0.9) of matched representative transcripts ^90^. Genes that did not map in MANE Select were next assessed for predicted consequences respective mutations have according to the Sequence Ontology ^91^. The aberrant transcript having the highest predicted impact was selected. The longest transcript was chosen when the same impact was annotated by Sequence Ontology for several transcripts representing the same gene. Genes in the ACCC CRC dataset with SVRACAS prioritized mutations were mapped to a curated list of CRC cancer driver genes in the Integrative OncoGenomics database (IntOGen) ^30^ (https://www.intogen.org, v1-2020).

### Identification and annotation of Copy Number Alterations

Copy Number Alterations (CNAs) were inferred from the whole exome sequencing data using a customized workflow (Supplementary File 1). Briefly, CNVkit (version 0.9.3) ^92^ was utilized with default parameters to infer discrete copy number segments. Next, tumor cell content and ploidy were estimated using a method adapted from ACEseq ^93^. Absolute copy numbers and allele-specific copy numbers as well as the decrease in heterozygosity (DH) ^94^ were estimated segment-wise, for each possible combination of Tumor cell content and ploidy.

### RNA-Sequencing and analysis

RNAseq libraries were prepared using the TruSeq Stranded library protocol (Illumina) and sequencing was done with a NovaSeq 6000 (Paired-end 100 bp S1) in the DKFZ Sequencing Core Facility to collect a median of 75 million reads/sample (spread: 65-99 million). Processing of RNA-Seq data was performed using a customized RNAseq Workflow as a part of the DKFZ – ODCF Roddy pipeline (https://github.com/DKFZ-ODCF/RNAseqWorkflow, version 1.3.0). After initial QC, the RNAseq raw reads were aligned to a customized hg19 version of the 1000 genomes reference genome (1KGRef_PhiX): (ftp://ftp.1000genomes.ebi.ac.uk/vol1/ftp/technical/reference/phase2_reference_assembly_sequence/hs37d5.fa.gz), by a 2 pass alignment using STAR aligner ^95^ (version 2.5.3a) and gencode 19 gene models. PhiX-genome sequence was added to the reference as an additional contig to remove spike-in sequences that had been included to find potential cross-contaminations and sample swaps. Additional QC after alignment was performed using samtools *flagstat* command and the RNA-SeQC tool ^96^ (version 1.1.8). The featureCounts function ^97^ as part of the Subread software package ^98^ (version 1.5.1) was used to determine the respective raw counts per gene, using gencode 19 gene models. Downstream RNASeq analysis was performed with R (R version 4.1.0)/Bioconductor software. The R package EnsDb.Hsapiens.v75 was utilized to annotate gene symbols. Non-specific intensity filtering was performed to exclude genes with less than 10 read counts in less than five samples. TMM normalization was conducted using the edgeR R package ^99^ (version 3.36.0), followed by incorporating sample weights and increase statistical power using the Voom function. Finally, the R package immunedeconv ^45^ (version 2.1.0) was utilized to estimate the composition and density of immune and stromal infiltration, using the non-log-transformed Transcripts Per Kilobase Million (TPM) values as input.

### TCGA and CPTAC multi-OMICs data

Public data from two CRC cohorts were downloaded from the cBioPortal multi-omics cancer portal (https://www.cbioportal.org/) i.e., the PanCancer Atlas TCGA Colorectal adenocarcinoma ^11^ and the CPTAC Prospective Colon ^100, 101^ datasets. For both datasets, only samples were processed that had been profiled for SNVs, CNAs, gene expression (RNAseq). Phosphoproteomic data were available only from CPTAC. SNVs were retrieved and data analysis was performed using the R package maftools ^102^ (version 2.10.0). The significance of differential mutational frequencies between the MSS and MSI tumors within TCGA and CPTAC cohorts was assessed using Fisher’s exact test based on a 2×2 contingency table (adjusted p-value ≤0.1, minimum number of samples harboring mutated genes ≥4), exploiting the mafCompare function of maftools.

### Inference of microsatellite instability status in the CRC datasets

We utilized the MSI MANTIS ^103^ estimators of microsatellite instability provided with the TCGA pan-cancer dataset, using the following categorization: A TCGA tumor was classified as MSI if the respective value from MSI MANTIS was higher than 0.6, while tumors with relative values lower than 0.4 were classified as MSS. Any tumors with relative values between 0.4 & 0.6 were denoted as “Undetermined” and excluded from further analysis. Information about MSI-status of respective samples was available also for the CPTAC dataset ^101^ and utilized as provided. Then, the PREMSIm R package ^23^ (version 1.0) was employed to infer the microsatellite status based RNA-sequencing data from all three datasets (TCGA, CPTAC, ACCC), using non-log-transformed Transcripts Per Kilobase Million values from RNA-sequencing. MSI-L and MSS tumors were grouped together in all datasets and the term MSS used for this group to distinguish from MSI tumors.

### Functional enrichment analysis

Functional Enrichment analysis was performed using the R packages clusterProfiler ^104^ (version 4.2.2), and ReactomePA ^105^ (version 1.38.0) was used for overrepresentation analysis based on REACTOME molecular pathways. ssGSEA ^44^ (https://github.com/broadinstitute/ssGSEA2.0) was implemented based on the ssGSEA projection methodology ^106^. The hallmark signatures file (version 7.5.1) was used in gmt format from the respective Molecular Signatures Database (MSigDB) ^107^ gene sets (https://www.gsea-msigdb.org/gsea/msigdb/). The script ssgsea-cli.R (https://github.com/broadinstitute/ssGSEA2.0) was run in a Windows 10 x64 (build 19044) environment using command line and the following arguments: *[-i ACCC.CRC.RNASeq.VoomCPM.TMM.Filtered –o ACCC –n none –d. /ssGSEA2.0-master/db/msigdb/h.all.v7.5.1.symbols.gmt –w 0.75 –c z.score –p 1000 –m 10]*. Protein associations were retrieved and visualized using STRING ^36^ (https://string-db.org<u>, accessed 2023/12/04</u>), and K-means clustering was performed to identify functional protein networks.

### Pathway, TF and Kinase activity inference analyses

The upper-quartile of processed and log2 transformed RSEM estimated counts from CPTAC RNAseq data were normalized using the function *normalizeQuantiles* from the limma R package ^108^ (version 3.50.0). Then, a non-specific intensity filtering procedure was implemented to remove features that were not expressed at least in the group with the lowest number of samples. The same preprocessing approach was applied for the TCGA dataset, with two additional steps: as the estimated raw RSEM counts had been batch effect-corrected, many features assigned by NA values were removed. Then, the data were log2-transformed if the maximum value was >50. Differential expression analysis in both datasets (MSI vs MSS) was applied with the limma R package ^108^.

The PROGENy (Pathway RespOnsive GENes R package ^37^ (version 1.16.0) was implemented to identify putative differentially activated molecular pathways, based on the derived differential moderated statistics extracted from the limma pipeline (i.e., top 100 most responsive genes for each pathway). PROGENy infers the activity of 14 major signaling pathways, based on the exploitation of consensus gene signatures and estimated from a large compendium of perturbation experiments ^37^. The DoRothEA-decoupleR pipeline ^109^ (version 1.6.0) was implemented to elucidate differentially activated regulatory networks. Relative activities were inferred for every transcription factor (TF) from the deregulated expression of its respective target genes, using the normalized weighted mean statistical metric from the decoupleR R package ^110^ (version 2.1.6). Customized barplots were created using the R package ggplot2 ^111^ (version 3.3.5) for the visualization of differentially altered TFs and pathways.

Phosphoproteomics data from the CPTAC cohort were processed and analyzed to infer differentially activated signaling cascades between MSS and MSI colon tumors. To this end, normalized phosphosite ratios were used, representing scaled intensity values. Duplicated entries were removed and any phosphosites were excluded that had NA values across more than 85% of the total samples. Differential protein phosphorylation analysis was performed using the limma R pipeline (version 3.50.0). Omnipath-decoupleR ^110^ and Post-translational Modification Set Enrichment Analysis (PTM-SEA) ^112^ with the PTMSigDB signaling pathways’ collection (version 2.0.0) were implemented to determine differentially activated kinases and signaling pathways, respectively. The Omnipath R package ^113^ (version 3.2.8) was utilized to retrieve the prior-knowledge interactions composed by kinase-target relationships, whereas the R package decoupleR ^110^ (version 2.1.6) was used to estimate the relative biological activities by application of statistical methods (normalized weighted mean).

### Patient clustering

Unsupervised hierarchical clustering on principal components (HCPC) method was implemented, based on the R packages FactoMineR ^114^ (version 2.8) and Factoshiny (http://factominer.free.fr/graphs/factoshiny.html, version 2.4). The selected scaled pathway and TF activities per patient were used as input for the *PCA* function. Then, hierarchical clustering on the first 8 components (accounting ∼95% of retained information) was performed using the *HCPC* (kk=Inf, max=10, nb.clust = –1, consol = TRUE) function. Plots were created using the ComplexHeatmap R package. Patients were re-ordered within each cluster according to their microsatellite instability status and selected mutational groups, namely BRAF_mut, KRAS_mut, NRAS_mut, GNAS_mut, RAS_RAF_GNAS_mut, and RAS_RAF_GNAS_wt.

### Identification of actionable alterations for construction of personalized cancer patient maps

Aiming to identify actionable vulnerabilities at the single patient resolution within the ACCC-CRC cohort, the qualified somatic variants from the SVRACAS scoring scheme were integrated with clinical implications from the OncoKB precision oncology database ^29^. Variant oncogenic effect annotation was applied to keep only variants characterized as oncogenic or likely oncogenic. Therapeutically actionable variants in any cancer type were assigned with an OncoKB Therapeutic Level of Evidence (https://www.oncokb.org/therapeutic-levels, accessed 2022/12/21). Genes carrying actionable variants were assigned to the treatment baskets of the National Center for Tumor Diseases/German Cancer Consortium (NCT/DKTK) MASTER (Molecularly Aided Stratification for Tumor Eradication Research) program, that are based on a set of 472 genes which participate in biological processes and cellular pathways with established therapeutic recommendations ^51^.

### Reporting summary

Further information on experimental design is available in the Nature Research Reporting Summary linked to this article.

## Data availability

All sequencing data supporting the findings of this study have been uploaded to the European Genome-Phenome archive (http://www.ebi.ac.uk/ega/), and are available with controlled access of a DAC via the accession numbers EGAD00001011262 and EGAD00001011263. All computational code, including R scripts, quarto documents and further information to reproduce the complete bioinformatics analyses, are available on Zenodo (https://zenodo.org/doi/10.5281/zenodo.10959699) and Github (https://github.com/Jasonmbg/ACCC-CRC, https://github.com/Jasonmbg/TCGAPanCan-CRC and https://github.com/Jasonmbg/CPTAC-Colon) repositories for open-access.

## Acknowledgements

We thank the teams of the DKFZ High-Throughput Sequencing and the OMICS IT Core Facilities for performing excellent services. We thank Julio Saez-Rodriguez and Aurelien Dugourd (Heidelberg University), and Marcos Díaz-Gay (University of California San Diego) for support concerning the implementation of pathway/TF activities, and COSMIC signature analysis, respectively. This study was supported by the Helmholtz European Partnering Program “Athens Comprehensive Cancer Center (ACCC)” in the course of a strategic collaboration between the National Hellenic Research Foundation (NHRF), the G. Gennimatas Hospital, the Medical School of the University of Athens, and the German Cancer Research Center (DKFZ).

## Author contributions

G.N.Z. **clinically assessed or operated patients and coordinated sample acquisition**. S.T., G.N.Z., V.K., S.R, Ko.P, and A.P. **collected and processed specimens**. S.T., S.Wo and S.Wi. **oversaw sequencing of the samples**. E-I.V., K.V., G.W., R.K., Ky.P., and O.P. **carried out the bioinformatics analyses and processed data**. E-I.V., K.V., J.B., O.P. and S.Wi. **analyzed and interpreted data, and wrote the manuscript**. E-I.V., A.D-S., G.N.Z., A.P. and S.Wi. **conceived the study**. **All authors participated in discussions and interpretation of the data and results**.

## Additional Information

Correspondence should be addressed to Efstathios-Iason Vlachavas, Olga Papadodima or Stefan Wiemann.

