## Supplementary figures and images for "Molecular and functional profiling unravels targetable vulnerabilities in colorectal cancer"

### Supplementary Figure 1

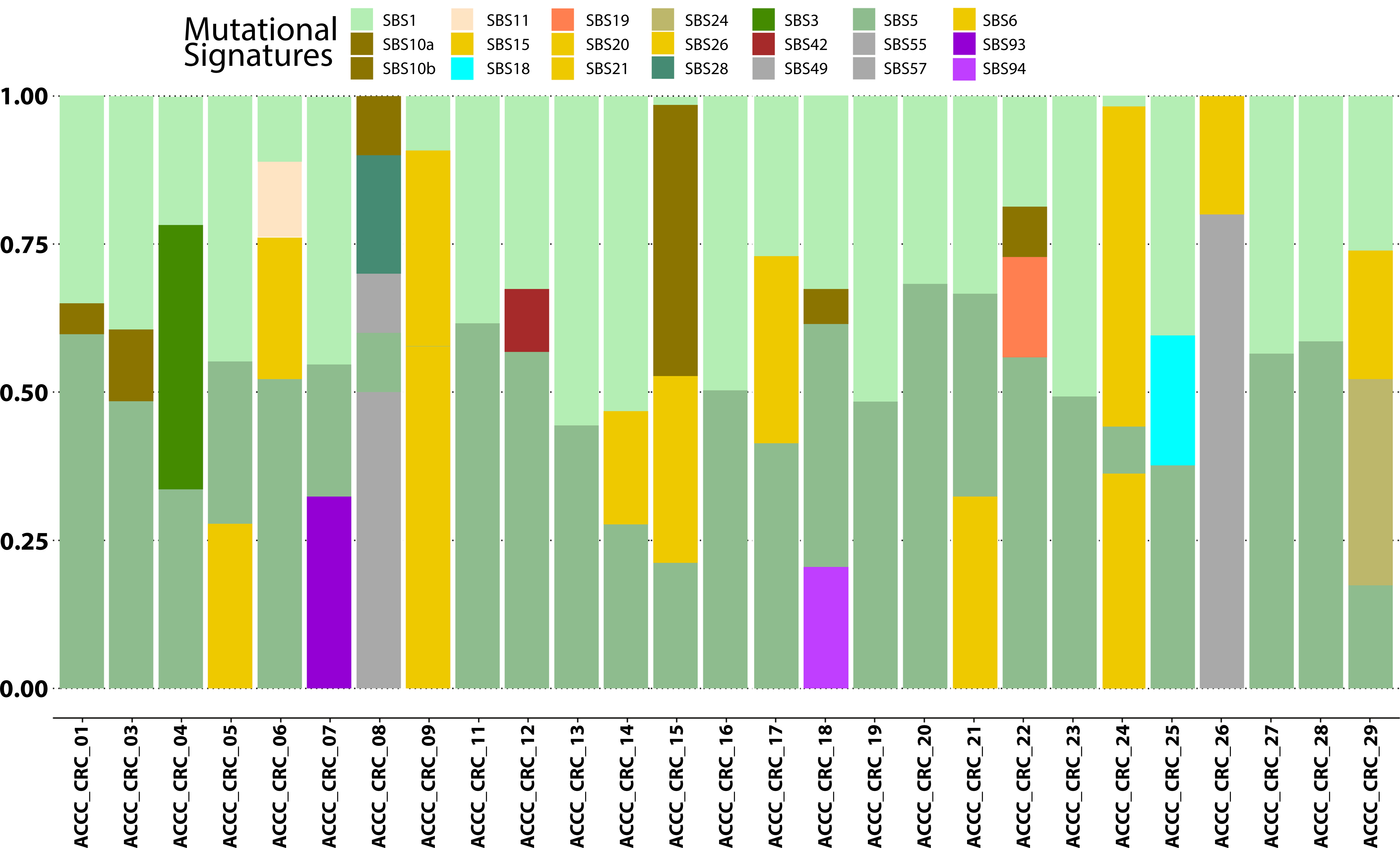

### Supplementary Figure 2

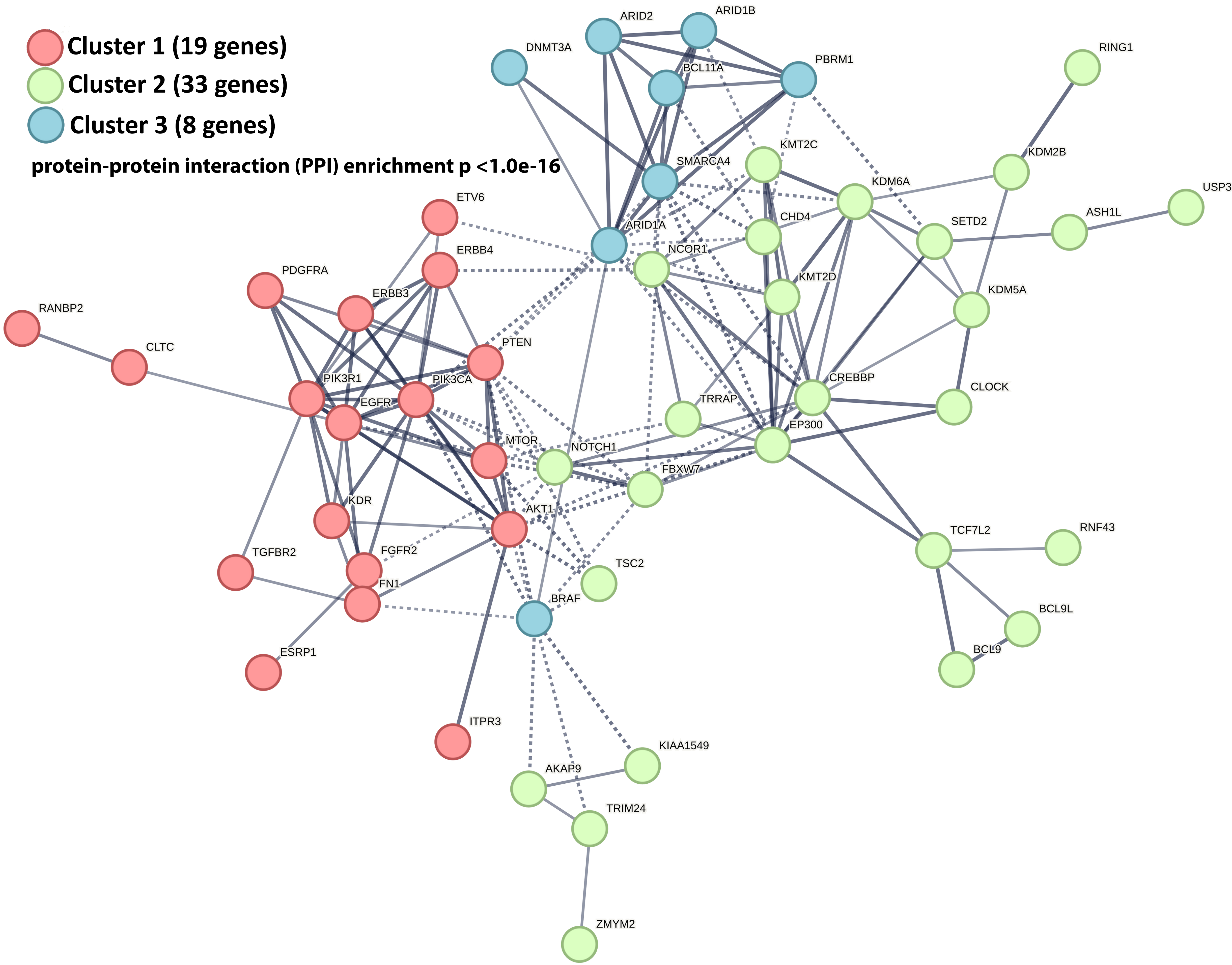

### Supplementary Figure 3

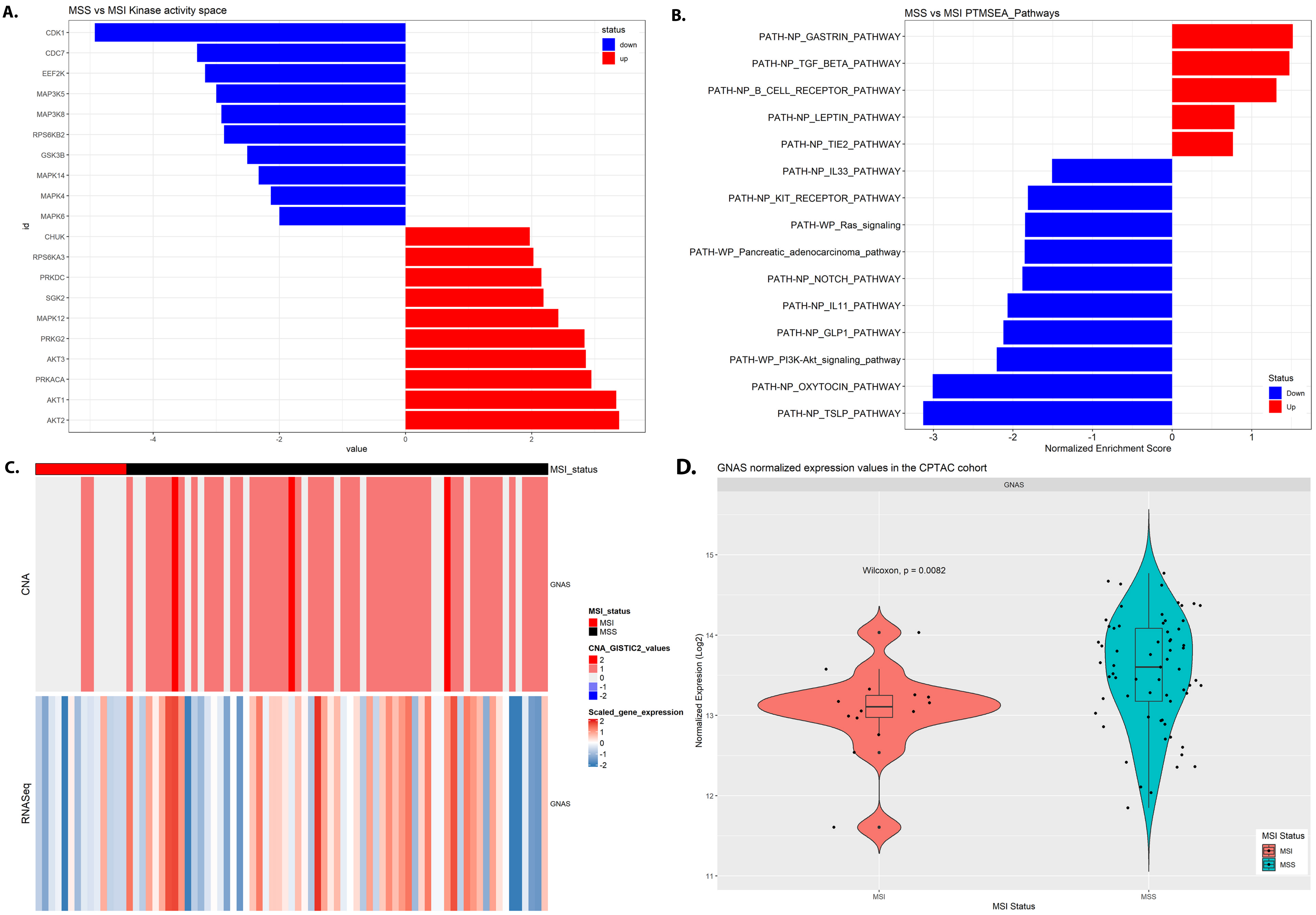
